# The genomic trajectory of ovarian high grade serous carcinoma is determined in STIC lesions

**DOI:** 10.1101/2024.03.11.584384

**Authors:** Z. Cheng, D.P. Ennis, B. Lu, H.B. Mirza, C. Sokota, B. Kaur, N. Singh, O. Le Saux, G. Russo, G. Giannone, L.A. Tookman, J. Krell, C. Barnes, J. McDermott, I.A. McNeish

**Affiliations:** Ovarian Cancer Action Research Centre, Department of Surgery and Cancer, Imperial College London, London, UK; Cancer Research UK Cambridge Institute, University of Cambridge, Li Ka Shing Centre, Cambridge, UK; Department of Cell and Developmental Biology, University College London, UK; Department of Cellular Pathology, Imperial College Healthcare NHS Trust, London, UK; Department of Pathology, Barts Healthcare NHS Trust, London, UK

**Keywords:** High grade serous carcinoma, STIC, evolution, whole genome duplication

## Abstract

Ovarian high-grade serous carcinoma (HGSC) originates in the fallopian tube, with secretory cells carrying a *TP53* mutation, known as ‘p53 signatures’, identified as potential precursors. p53 signatures evolve into serous tubal intraepithelial carcinomas (STIC) lesions, which, in turn, progress into invasive HGSC that readily spread to the ovary and disseminate around the peritoneal cavity. We recently investigated the genomic landscape of early- and late-stage HGSC and found higher ploidy in late-stage (median 3.1) than early-stage (median 2.0) samples. Here, to explore whether the high ploidy and possible whole genome duplication observed in late-stage disease are determined early in the evolution of HGSC, we analysed archival formalin-fixed paraffin-embedded samples (FFPE) from five HGSC patients. p53 signatures and STIC lesions were laser-capture microdissected and sequenced using shallow whole genome sequencing (sWGS), while invasive ovarian/fallopian tube and metastatic carcinoma samples underwent macrodissection and were profiled using both sWGS and targeted next generation sequencing. Results showed highly similar patterns of global copy number change between STIC lesions and invasive carcinoma samples within each patient. Ploidy changes were evident in STIC lesions, but not p53 signatures, and there was strong correlation between ploidy in STIC lesions and invasive ovarian/fallopian tube and metastatic samples in each patient. The reconstruction of sample phylogeny for each patient from relative copy number indicated that high ploidy, when present, occurred early in the evolution of HGSC, which was further validated by copy number signatures in ovarian and metastatic tumours. These findings suggest that aberrant ploidy, suggestive of whole genome duplication, arises early in HGSC, and is detected in STIC lesions, implying that the trajectory of HGSC may be determined at the earliest stages of tumour development.

## Introduction

High grade serous carcinoma (HGSC) is the commonest subtype of ovarian cancer, but its cell of origin remained unclear until recently. In 2001, Piek et al first described preneoplastic lesions in the normal fallopian tube of women at high familial risk of HGSC undergoing risk-reducing surgery [1]. The subsequent development of Sectioning and Extensively Examining the FIMbria (SEE-FIM) [2] also allowed identification of non-proliferating secretory-type cells with aberrant p53 staining, so-called “p53 signatures”, as potential precursor lesions in the secretory epithelium of the fallopian tube fimbria [3]. p53 signatures are thought to transform into serous tubal intraepithelial carcinomas (STIC) via serous tubal intraepithelial lesions (STIL) [4].

STIC lesions have a clonal relationship with established HGSC based on shared *TP53* mutations [5], and whole-exome sequencing (WES) has confirmed that p53 signatures and STICs serve as precursors of ovarian carcinoma [6,7]. Mutation rates, mutational signatures and somatic copy number alterations (sCNA) are consistent between STIC and HGSC but unique to each patient. These alterations also appear consistent between anatomic sites within each patient, suggesting the biological processes underlying genomic instability during the HGSC evolution are persistent and stable.

We recently showed that the HGSC genome is remarkably stable between diagnosis and relapse and that acquired chemotherapy resistance does not select for common copy number drivers [8]. However, it remains unclear whether, and to what extent, genome-wide changes alter during the evolution of HGSC from initiation until the time of diagnosis. In our previous study [9], we compared the genomes of early-(stage I/IIA) and late-stage (stage IIIC/IV) HGSC, revealing no significant differences in the rates of somatic mutations between early- and late-stage samples, and no cohort-specific sCNA. However, high ploidy, suggestive of whole genome duplication (WGD), was observed frequently in late-stage disease but rarely in early-stage. However, it was unclear whether the features observed in late-stage cases were simply time-related markers of evolutionary fitness or whether they arose early during carcinogenesis as potential drivers of poor prognosis and metastatic dissemination.

To address this question, we have undertaken genomic analysis of a cohort of patients with HGSC in which we were able to identify p53 signature, STIC and invasive carcinoma within the each case. Our data suggest that features associated with advanced disease, including high ploidy, can be observed in STIC lesions. Moreover, there is remarkable consistency of ploidy and sCNA between lesions within individual patients, suggesting that the genomic landscape of HGSC is determined at the earliest stages of carcinogenesis.

## Materials and Methods

### Study conduct and patient samples

The samples were obtained under the authority of Imperial College Healthcare Tissue Bank (HTA licence 12275; REC approval number 17/WA/0161; Project ID R18060). All patients gave written consent. Samples were reviewed by expert gynaecological pathologists.

### Laser capture microdissection and DNA extraction

p53 signatures, STIL and STIC lesions were identified as previously [4] (see Supplementary Methods). Stroma, normal fallopian tube epithelium, p53 signature and STIL/STIC lesions were laser-capture microdissected (Carl Zeiss, Germany). Invasive carcinoma in the ovary and metastatic sites were macrodissected and DNA extracted from 10 × 10 μm sections using QIAmp DNA FFPE Tissue Kit (Qiagen, UK).

### Sequencing

Microdissected DNA samples were repaired by NEBNext FFPE DNA Repair Mix (M6630) and the NEBNext Ultra II DNA library Prep Kit (E7645) was used for whole genome library preparation. Shallow whole genome sequencing (sWGS) was performed on a HiSeq4000 system (Illumina Cambridge, UK), using paired-end 150 bp protocols. Analysis of *PTEN, KRAS, RB1, BRCA2, RAD51B, FANCM, PALB2, RAD51D, TP53, RAD51C, BRIP1, CDK12, NF1, BRCA1, BARD1, PIK3CA* was performed using a custom Ampliseq panel on a HiSeq4000 system (Illumina, Cambridge, UK), using paired-end 150 bp protocols [9]. All sequencing data are available via the European Genome-phenome Archive at the European Bioinformatics Institute (https://ega-archive.org) with accession number EGAS00001005567.

### Mutation calling

FASTQ files from AmpliSeq were aligned to reference human genome hg19 using Burrows-Wheeler Alignment (BWA-MEM) [10] and pre-processed using samtools and Picard to generate sorted BAM files [11]. Somatic mutations were called using Mutect2 (GATK4.1.4.1) [12] and Strelka [13] as previously [9].

### Absolute copy number and copy number signature calling

sWGS reads were aligned to reference human genome hg19. QDNAseq [14] and CGHcall [15] were utilised to obtain relative copy numbers in bins of 500 kb. Consecutive bins with the same copy numbers were merged across all samples into segments. In samples where laser capture yielded insufficient materials for both Ampliseq and sWGS, we derived the absolute copy number by adjusting the relative copy number based on the ploidy of each sample. Focal gene changes were then identified based on inferred absolute copy number. We defined gain as total copy number >2.5 and loss as a total copy number ≤1.5 [16]. Where STIL and STIC lesions were observed in the same patient, samples were included in the same classification group (‘STIC’) for copy number and ploidy analyses.

Where there was sufficient DNA for both Ampliseq and sWGS, we utilised sWGS-absoluteCN (swgs) to infer absolute copy number [8] and then used this to calculate CN signatures [17]. CN signatures were compared with those from our previous early stage and late-stage samples using cosine similarity function from R package lsa [18] on pairwise analysis of signatures-by-samples.

### Statistical analyses

Statistical analyses were conducted using Prism (v10.1.1, GraphPad). For comparing means between two groups, *t*-tests were employed for populations with a normal distribution, and Mann– Whitney tests were utilized for nonparametric distributions. Cluster distribution by cosine similarity was compared by one-way ANOVA. Correlation between ploidy in STIC and invasive samples and between *TP53* absolute copy number and ploidy were visualized using a scatterplot and statistically tested using Pearson’s (normally distributed) correlation test. Throughout, p<0.05 is considered significant.

## Results

### STIC clinical cohort

We identified five patients diagnosed with HGSC who had undergone surgery, including bilateral salpingo-oophorectomy, whose archival pathology samples were available at Imperial College Healthcare NHS Trust, London, UK and where p53 signatures, STIC lesions and invasive carcinoma were all detectable in each patient’s samples. A summary of the workflow is presented in Figure 1A, and the clinical information is presented in Supplementary Table 1. Following expert pathology review (CS, BK, NS, JM), 5 µm sections from each formalin-fixed, paraffin-embedded fallopian tube sample underwent immunohistochemistry (IHC) staining for p53 and Ki67 as well as H&E staining to identify STIL and STIC lesions and p53 signatures (Supplementary Fig. 1-5) using validated algorithms [4]. Subsequently, samples from invasive tubo-ovarian carcinomas (and metastatic tumours where present) were collected from the same patients. This allowed us to evaluate p53 signature lesions from four patients (STIC_0005, STIC_0012, STIC_0013, STIC_0014), STIC lesions and invasive carcinoma from the fallopian tube and/or ovary in all five patients, and omental, peritoneum, transverse colon, or aorto-caval lymph node metastases in four patients (STIC_0001, STIC_0005, STIC_0012, STIC_0014) (Supplementary Table 2). STILs were identified in all five patients but were included in the same classification as STIC lesions for genomic analyses. Germline *BRCA1/2* mutation data were available for two of the patients, neither of whom had a pathogenic germline alteration.

**Figure 1.**
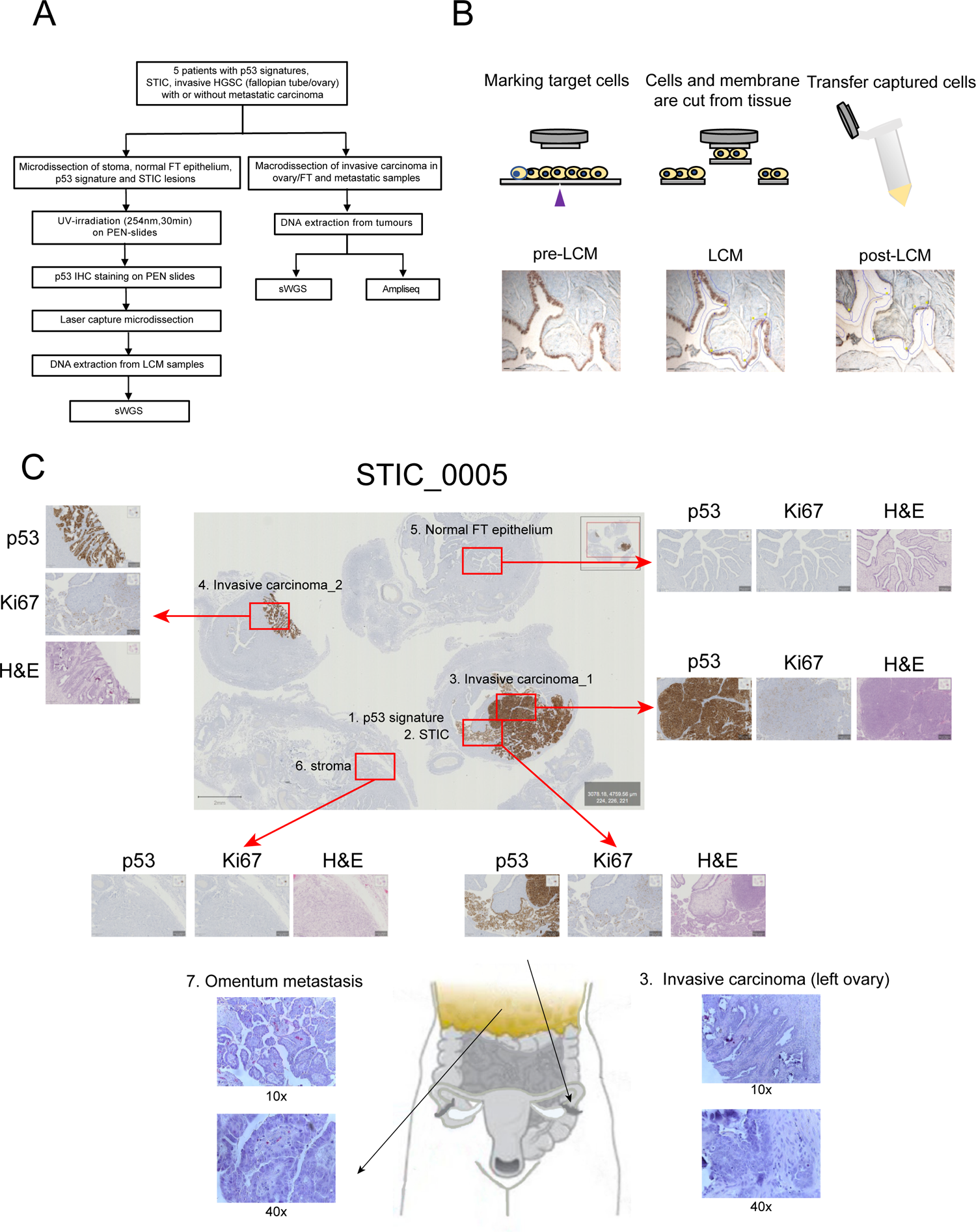
Workflow for identifying HGSC STIC cohort A. Workflow of sample isolation and next-generation sequencing analyses. For p53 signature and STIC samples, slides were stained by immunohistochemical staining of p53. Tumour samples were microdissected for sWGS analysis. Next-generation sequencing analyses were performed for tumour specimens using both sWGS and Ampliseq sequencing for primary and metastasis carcinoma samples. B. The process of laser capture microdissection using the PALM Zeiss UV laser system is illustrated. PALM software was used to mark the target cells of interest, and a UV laser was then used to cut away these cells. The cells, along with the membrane, were then ejected against gravity and collected in an adhesive cap for downstream DNA extraction (as shown in the figures above). A representative figure demonstrates the laser capture microdissection of the STIC lesion before and after the process, with the STIC lesion stained in p53 immunochemistry (as shown in the bottom figures). C. Representative p53, Ki67 and H&E images of patient STIC_0005 with stroma, normal fallopian tube epithelium, p53 signature, STIC, and invasive ovarian tumour, metastasis sites. The samples in this patient are: 1. p53 signature; 2. STIC; 3. invasive tumour_1; 4. invasive tumour_2; 5. Normal fallopian tube epithelium; 6. Stroma; 7. Metastasis in omentum.

To isolate DNA from p53 signatures and STIC lesions, laser capture microdissection (LCM) was employed (Fig. 1B). Importantly, we collected DNA from individual lesions separately when multiple STIC lesions were identified within the same block. Normal fallopian tube stroma and epithelium were microdissected from the same slide as the p53 signature and STIC lesions to serve as controls. For invasive carcinoma in the fallopian tube/ovary and for all metastasis samples, macrodissection was performed from FFPE blocks after H&E staining. Examples of sample selection are shown in Fig. 1C and Supplementary Fig. 6–9.

### The genomic landscape of invasive carcinoma samples

The DNA yield from STIC_0013 was insufficient for panel sequencing, but targeted next-generation sequencing of macro-dissected invasive carcinoma and metastatic samples from the remaining four patients revealed mutations in *TP53* in 10/11 samples (91%). The one sample in which a *TP53* mutation could not be identified, from patient STIC_0005, was collected following neoadjuvant chemotherapy and had low tumour cellularity. However, p53 IHC (Supplementary Fig. 1) on this sample showed intense nuclear staining in keeping with a *TP53* missense mutation [19].

No somatic mutations were identified in *BRCA1* or *BRCA2*. However, we identified a missense mutation in *BRIP1* in all three samples from patient STIC_0012 and a truncating mutation in *NF1* in patient STIC_0001 (Fig. 2A).

**Figure 2.**
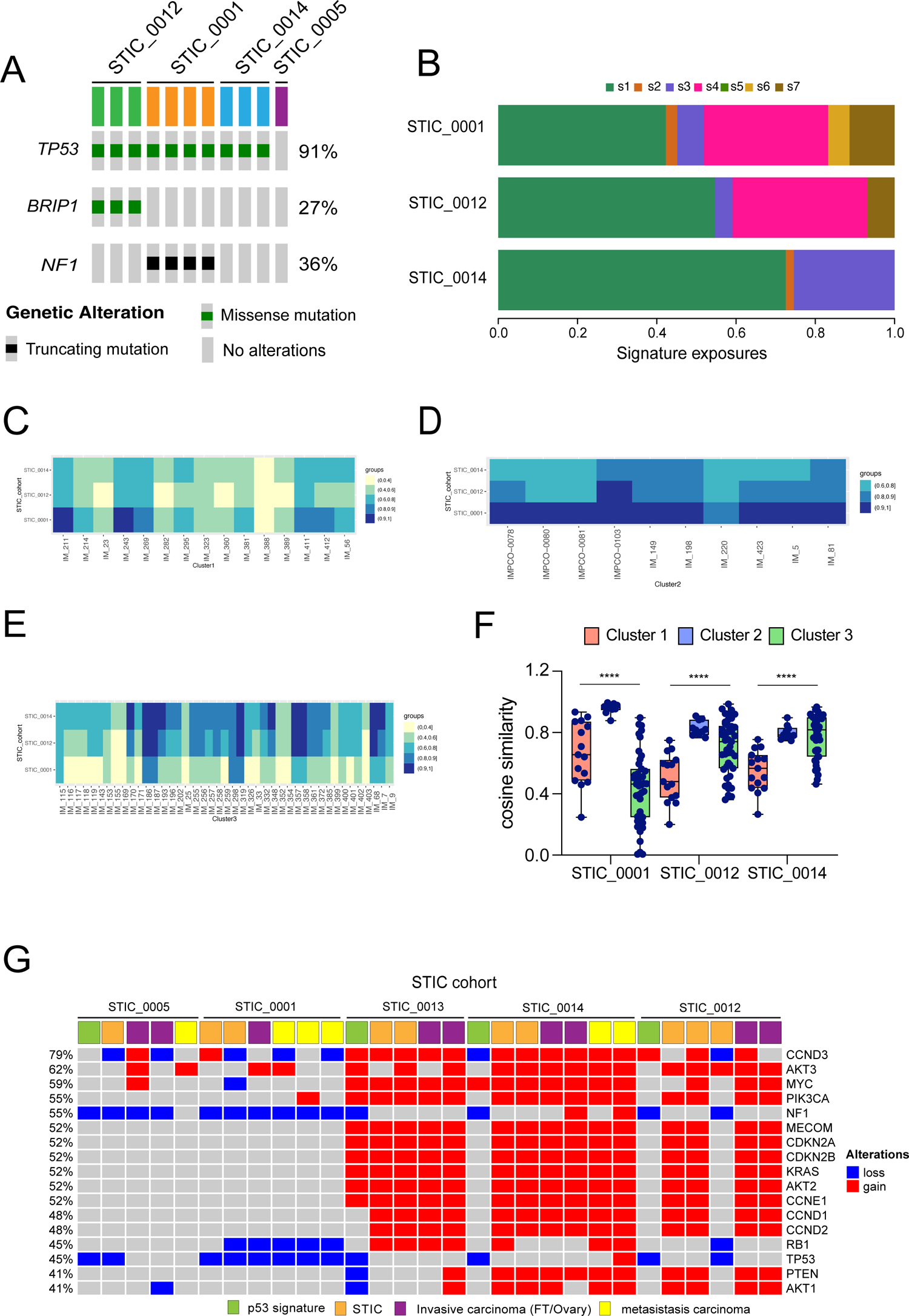
The genomic landscape of invasive carcinoma samples A. Somatic mutations for each patient in STIC cohorts. B. Copy number signature exposures in the STIC cohort (n=3). C–F. Cosine similarity comparison between STIC cohort and early/late-stage cohort by copy number cluster (p<0.05). G. Focal amplifications and deletions estimation in 17 genes of interest, determined by relative copy adjusted to ploidy in the STIC cohort.

Samples from patients STIC_0001, _0012 and _0014 yielded DNA of sufficient quantity for assessment of copy number signatures [8,17] (Fig. 2B). We calculated cosine similarities between these samples and the prognostic three copy number signature clusters that we previously described [9] (Fig. 2C–E): cluster 1, associated with poor outcome, primarily represented genomes with high copy number (CN) signature 1 exposure, cluster 3 displayed the highest CN signature 4 exposure and was associated with good outcome, whilst cluster 2 represented an intermediate state. Cluster distribution differed significantly for each patient (Fig. 2F; all p<0.0001). STIC_0001 showed the highest similarity with cluster 2 (>0.95), while STIC_0014 had highest similarity with cluster 3 (0.82, p<0.001). For STIC_0012, there was highest similarity for cluster 2 (0.82).

### The copy number landscape of p53 signatures and STIC lesions

We utilised shallow whole genome sequencing to analyse copy number alterations in matched tumour and normal specimens from all patients. However, identifying copy number alterations in p53 signatures and STIC lesions was challenging. Therefore, we developed experimental and bioinformatic approaches to detect copy number from microdissected tissue. These included optimised microdissection after immunohistochemical staining, improved DNA recovery and library construction from limited and stained tissue samples (see Methods). Since p53 signatures are extremely small, frequently containing no more than a few hundred cells in total [7], yielding very limited amounts of isolated DNA (less than a few ng), it was necessary to utilise different bin sizes to analyse sWGS data. After comparing the segment numbers, relative error, purity and ploidy results using the ACE package [20] (Supplementary Figure 10), we choose 500kb as the standard bin size for downstream analysis. To ensure the reliability of the results, we analysed the purity and ploidy results in all samples, especially in p53 signatures and STIC lesions, using three different pipelines: ACE [20], Rascal [https://github.com/crukci-bioinformatics/rascal] and ichorCNA [21] (see Supplementary Table 3). The final purity and ploidy results for each sample were determined by the average from the three pipelines.

After computing overlap of reconstructed ancestral copy number profiles with COSMIC genes in HGSC [22], we summarised focal amplification and deletions (Supplementary Table 5) across all samples (Fig. 2G). Among the 18 genes that are most frequently amplified or deleted in HGSC, we found striking consistency between STIC lesions and invasive carcinoma samples from the same patient. We also identified amplifications in *MECOM* (3q26) [23], *MYC* (8q24) [24] and *CCNE1* (19q12) [25] in both p53 signature and STIC lesions, as well as universal deletions of *TP53* and *NF1* on chr17p in p53 signatures. Given that *TP53* loss is related to initiation of chromosome instability [26] and may play an important role in transforming normal epithelium into p53 signature [27], these data strongly suggest that genomic instability appears very early in the evolution of HGSC.

Furthermore, using GISTIC 2.0 [28], we revealed a high degree of genomic instability as early as p53 signatures (Fig. 3A, 3B). We also identified statistically significant regions of aberration (Fig. 3C), such as loss at cytobands 6p and amplification at 6q in p53 signatures, and amplification peaks at 6q and deletion peaks at 4q and 8p in STIC lesions. These locations are consistent with the previous GISTIC analysis of STIC samples [6]. The overall genomic location of CNA in the STIC cohort was also highly concordant with TCGA analysis of HGSC, including frequent amplification of chromosomes 1q, 3q, 6p, 8q, 20, and deletion of 4q, 6q, 8p, 11q, 18q, 22q [29], suggesting again that focal chromosome changes occur very early in HGSC development.

**Figure 3.**
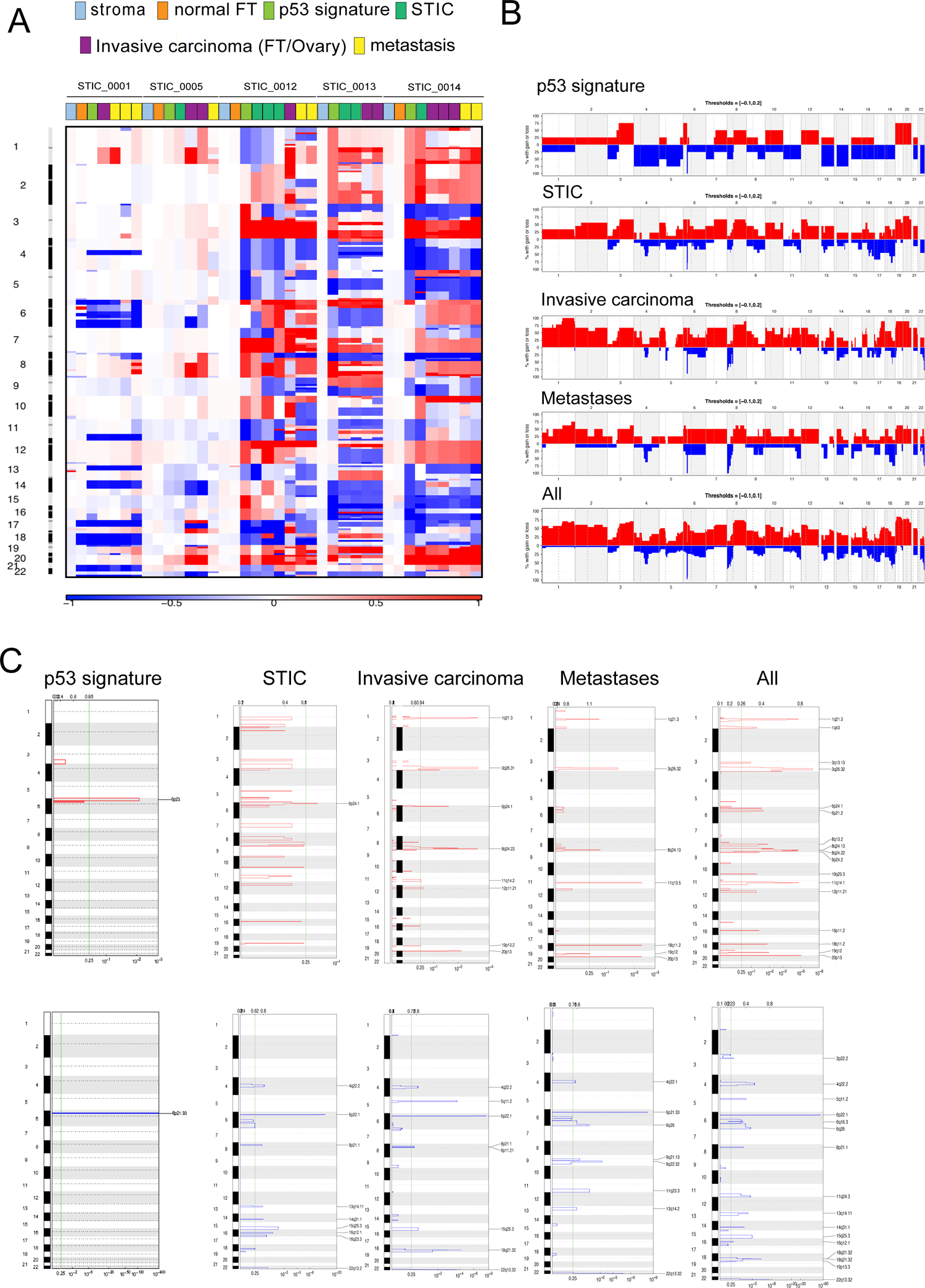
Analysis of genome-wide copy number alterations in the five patients. A. The plot of genomic aberrations of copy number alterations (CNA) across all patients and anatomic sites indicates a high degree of genomic instability in all regions. The chromosome numbers are marked on the margins, with amplifications in red and deletions in blue. B. The frequency of losses and gains in p53 signatures, STICs, FT/ovarian tumours, metastatic tumours, and overall samples are represented on a genome-wide scale, with gains marked in red and deletions in blue. The threshold for identification of these genetic alterations was set at [-0.1, 0.2]. C. The genome is depicted vertically from top to bottom, while the log-scaled GISTIC q-values at each locus are plotted horizontally from left to right. A green line is displayed to represent the significance threshold (q-value = 0.25).

### Ploidy changes with recurrent molecular alterations in the evolution of HGSC

Given our previous data on differences in ploidy between early and late stage HGSC, we next examined ploidy across samples (Fig. 4A). p53 signatures were diploid in all patients. However, ploidy changes were observed in STIC lesions. Interestingly, not all patients had the same ploidy changes, and the evolutionary trajectories were also different. In patient STIC_0001 and STIC_0005, where STIC lesions were diploid, the ovarian and metastasis sites were also diploid. By contrast, in STIC_0013 and _0014, STIC samples had high ploidy (>2.7), and high ploidies were also observed in the ovarian and metastatic samples. In the remaining case, STIC_0012 with three STIC lesions, two had higher ploidy (>2.5) and one was diploid, whilst the invasive carcinoma had high ploidy (carcinoma 2.5; metastatic samples 2.9, 3.0). Overall, there was a very strong correlation between ploidy in STIC lesions and that in matched invasive carcinoma and metastases (Fig. 4B). This strongly suggests that ploidy changes occur early in HGSC development and suggests that whole genome duplication could be a key driver in HGSC development [30].

**Figure 4.**
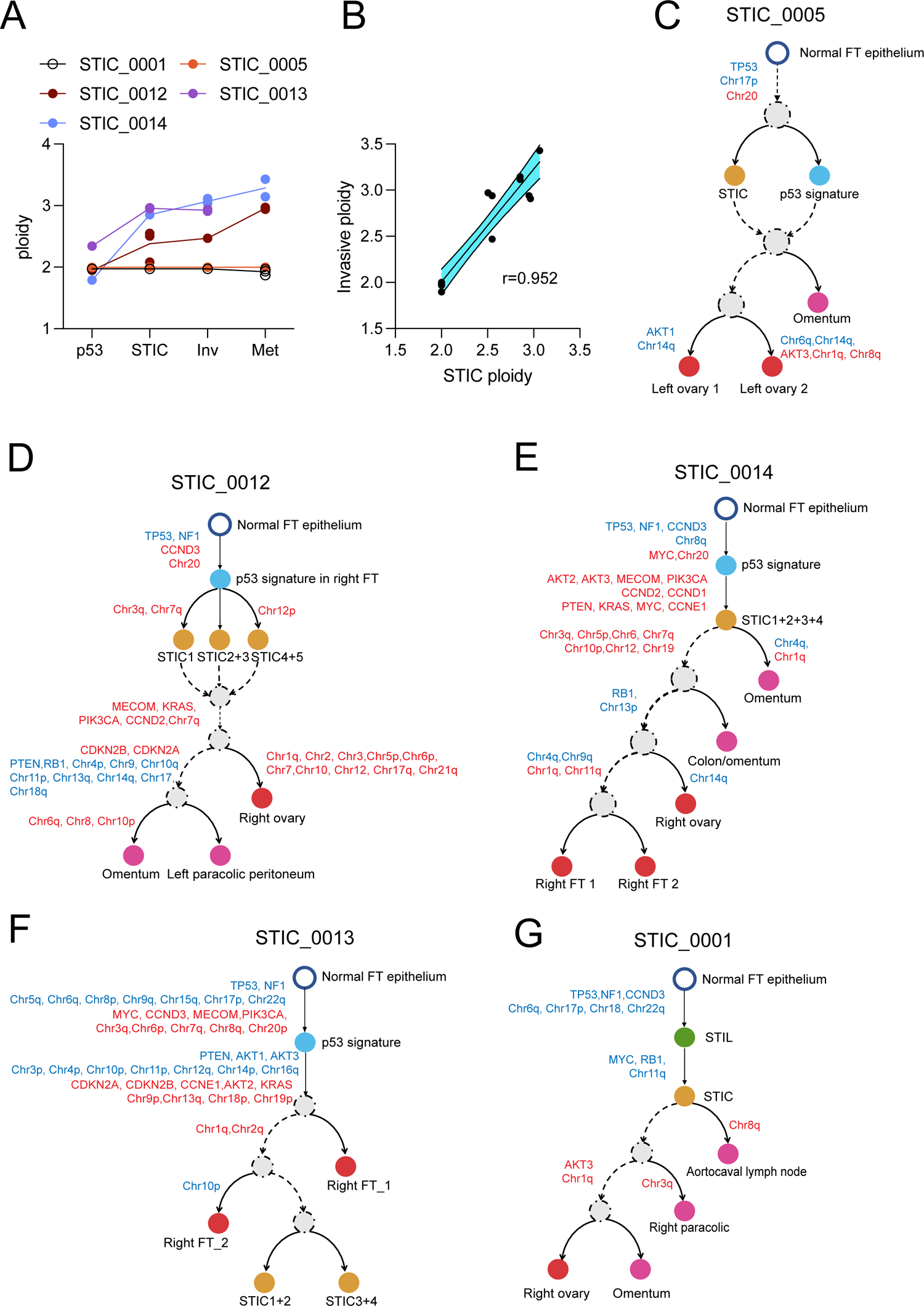
Copy number alteration phylogeny in STIC cohort A. The changes in ploidy during the evolution of five HGSC patients are depicted. B. Correlation between ploidy in STIC and invasive carcinoma samples within each patient. C–G. The evolutionary history of tumours in each patient is represented as a copy number hierarchy inferred from the genomic regions using the CNETML package and visualised as a tree structure with a root node corresponding to the normal fallopian tube epithelium. In all patients, loss of *TP53* is one of the earliest alterations and is present in all anatomic samples. Cosmic genes and chromosomes are gained or lost along the branches of the tree, and alterations are indicated at each node. Each node is labelled with the tumour samples harbouring all the upstream alterations and lacking any downstream alterations. Red indicates gene or chromosome gains, while blue indicates gene or chromosome losses.

### Unravelling the genomic evolutionary relationships in HGSC development

The conventional approach for analysing phylogenetic trees typically relies on somatic mutations or allele-specific copy numbers. However, due to the limited sample input, we applied CNETML [31], a new maximum likelihood method than can infer phylogenetic trees from relative copy number called from sWGS data, to determine the evolutionary trajectory of lesions for each patient (see Supplementary Methods). To visualise the evolutionary process of HGSC better, each phylogenetic tree was transformed into a hierarchy graph based on the inferred tree topology (Supplemental Fig. 11) and reconstructed ancestral copy numbers from known information on HGSC development and the quality of detected copy numbers (Fig. 4C–G).

Despite the background noise produced by FFPE artefacts and immunohistochemistry staining during LCM, the phylogenetic analysis of the evolutionary relationship provides evidence that nearly all alterations within the p53 signature/STIC lesions or their immediate precursors were shared by other lesions. This suggests strongly that they present the direct ancestral clone for the invasive carcinomas and metastatic tumours (Supplementary Fig. 11).

### *TP53* absolute copy number status is related to the ploidy changes

*TP53* mutations are considered to be the initiating event in high grade serous carcinoma development [26] and drive non-random patterns of chromosomal anomalies [32]. The rate of missense *TP53* mutations is approximately twice that of null [33], although there does not appear to be a relationship between mutation type and clinical outcome [34]. Here, we analysed relative copy number and adjusted to the estimated ploidy of each sample, allowing us to infer *TP53* absolute copy number.

*TP53* loss was universal in the p53 signatures (Fig. 5A), confirming that loss of the wild type *TP53* allele is observed in the earliest precursor lesions of HGSC. We also observed a strong correlation between *TP53* absolute copy number with ploidy across all samples (R=0.79, Fig. 5A). This analysis revealed two distinct groups of samples: those with one copy of *TP53*, in which ploidies were all low (<2.7), while the majority of those with two copies of *TP53* exhibited ploidies >2.7. This strongly suggests that whole-genome duplication occurs early after the transformative mutational events involving *TP53* and other cancer genes. However, *TP53* dysfunction is not an obligatory event for WGD [35,36]; indeed, the presence of wild-type p53 may be an absolute requirement for WGD that is driven by cyclin E1 [37]. Most LOH events are due to strict copy-loss (copy-loss LOH), where allelic loss occurs in the context of a decrease in gene copy number. However, copy-neutral LOH is also frequently observed, whereby an allele is lost but the number of gene copies remains the same, or even increases, due to chromosomal gain [38].

**Figure 5.**
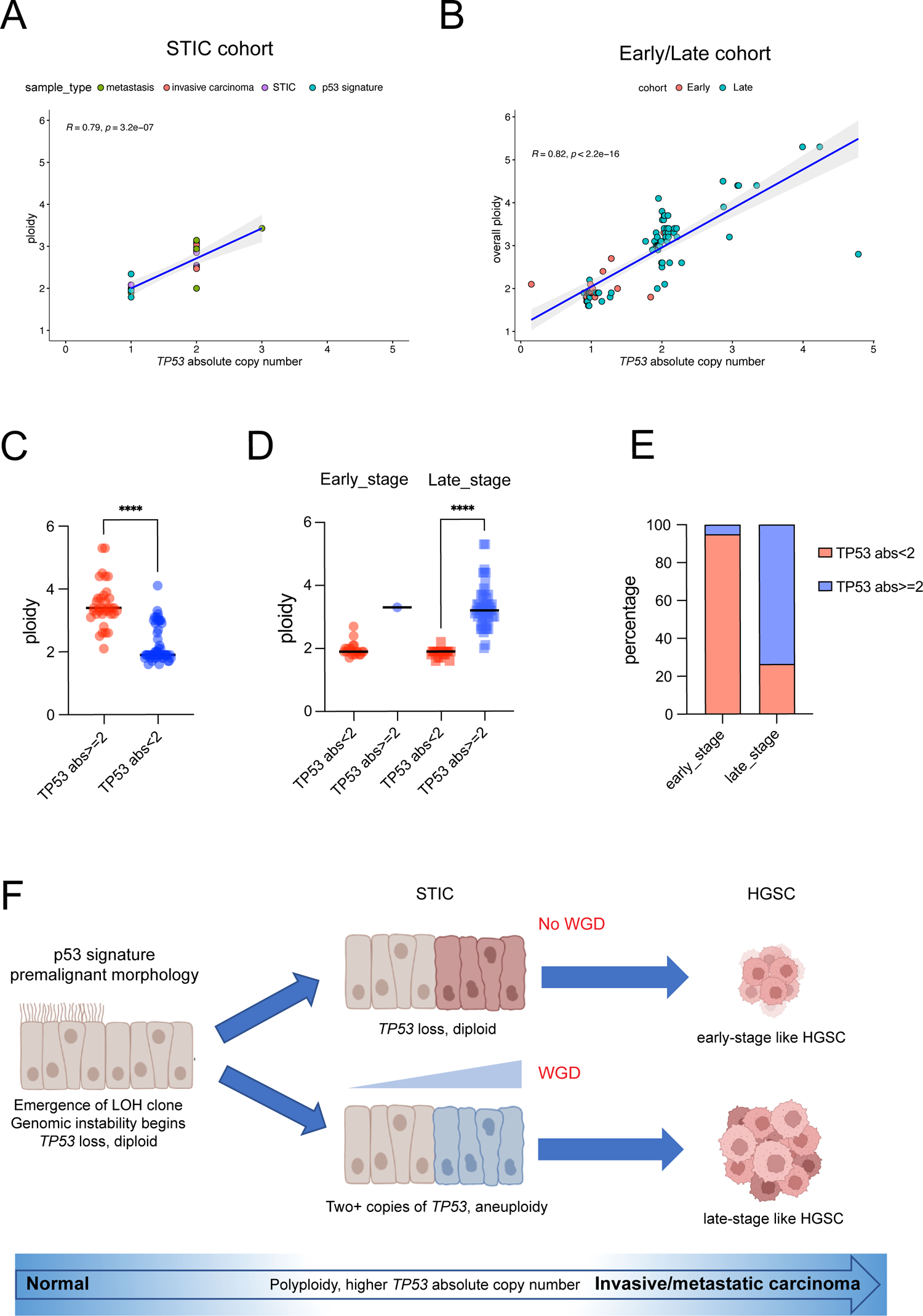
TP53 absolute copy number correlates with ploidy changes in the evolution of HGSC A. The correlation between *TP53* absolute copy number and ploidy in STIC cohort. B. The correlation between *TP53* absolute copy number and ploidy in combined early stage and late-stage cohort. C. Ploidy comparison between *TP53* absolute copy number (>=2 vs <2, p<0.05). D. *TP53* absolute copy number related to ploidy in early and late-stage cohort (p<0.05). E. Proportion of *TP53* absolute copy number distribution in early and late-stage cohort. F. The diagram depicts the gradual progression from p53 signature to STIC, and to ovarian or metastatic tumours, which exhibit *TP53* and ploidy changes.

To validate our observation further, we performed a combined analysis of all samples from our previous early and late-stage HGSC cohorts [9], again revealing a strong correlation between *TP53* absolute copy number and ploidy (Fig. 5B), whereby samples with two copies of *TP53* showed higher ploidy than those with a single copy of *TP53* (Fig. 5C). Interestingly, we did not observe any significant differences in ploidy between missense and non-missense *TP53* mutations in this larger cohort, indicating that changes in ploidy are not specifically associated with mutation type (Supplementary Fig. 12A, B). We also analysed our previous early and late-stage cohorts separately, demonstrating that the large majority of samples in the early-stage cohort had one copy of *TP53* and exhibited a diploid status (Fig. 5D), whilst the majority of late-stage samples had two (or more) copies of *TP53* and higher ploidies (Fig. 5E). These findings are consistent with the observations made in the current STIC cohort, suggesting that the distinction between early and late-stage features may be determined very early in the evolution of HGSC through the occurrence of WGD in the STIC lesions.

## Discussion

Tubo-ovarian high grade serous carcinoma is a disease of poor prognosis and extreme genomic complexity, marked by profound genomic instability and copy number alterations [29,39], that has largely failed to benefit from precision medicine approaches. Its origins at the distal end of the fallopian tube and early metastatic dissemination around the peritoneal cavity have made it challenging to examine the earliest stages of carcinogenesis. Here, we present an evolutionary analysis of multi-site HGSC samples in five patients. We employed phylogenetic tree analysis, as well as ploidy and *TP53* absolute copy number data, to investigate the evolutionary relationship between p53 signatures, STIC lesions and areas of invasive carcinoma on the ovary and distant metastatic sites. Our key finding is that profound copy number alteration is evident in STIC lesions, with ploidy change suggestive of WGD, detected as early as STIC lesions in some patients. Moreover, if high ploidy is detected in invasive carcinoma and metastatic samples, it is also detected in STIC lesions. This suggests strongly that the trajectory of HGSC is determined early in the carcinogenesis process, with prognosis potentially determined at the point of STIC emergence (Graphical representation in Fig. 5F).

Our findings extend previous research [7] and support the notion that ovarian cancer originates from the fallopian tube and undergoes a series of transitions from normal epithelium to p53 signatures and STIC lesions [40]. Genomic analysis revealed early chromosomal alterations occurring at the p53 signature and STIC stages, affecting key driver genes and pathways including *TP53*, cell cycle (*CCNE1, CCND1*), PI3K/RAS (*NF1, PTEN, PIK3CA, AKT*) and oncogenic signalling (*MYC*) [27]. Subsequently, the development of HGSC in the ovaries and more distant metastatic sites can be attributed to a seeding event originating from a primary tumour in the fallopian tube that already exhibits sequence and structural alterations in these driver genes. The recurrent allelic imbalances observed in chromosomes 1q, 3q, 4q, 6p, 8q, 22q indicate the potential involvement of additional genes and pathways in the development of HGSC [41,42].

Loss of p53 function is believed to be the initiating event in HGSC carcinogenesis. Certainly, *TP53* loss is a driver of subclonal karyotype alterations and initiation of copy number instability in fallopian tube epithelial cells [26]. However, whether this alone is sufficient to drive carcinogenesis is unclear. Certainly, in normal oesophagus, mutation of a single *TP53* allele was insufficient to allow tumorigenesis in the absence of LOH [43], whilst in pancreatic cancer mice models, single cell sequencing revealed four distinct and ordered phases of genomic instability following p53 inactivation, namely *Trp53* LOH, followed by accumulations of deletions, whole genome doubling and emergence of gains/amplifications, which correlated with tumour progression [32]. Our observation that allelic loss of *TP53* is universal in p53 signature lesions suggests that, by the time these lesions emerge, full (or near-full) transformation may already have occurred, in keeping with the classical Knudsen two-hit hypothesis [44]. To explain the differences in *TP53* absolute copy number observed in low and high ploidy STIC lesions, we hypothesise that there is either selective reduplication of the mutated *TP53* allele through copy number-neutral LOH, as is observed in early onset colorectal carcinoma [30], or whole genome duplication. Our data, showing high correlation between ploidy and *TP53* absolute copy number, suggest the latter. However, addressing this hypothesis fully would ideally require deeper whole genome sequencing to obtain B-allele frequency, although this would be technically highly challenging in small STIC lesions. Alternatively, fluorescent in-situ hybridisation (FISH) for *TP53* probes would allow absolutely CN quantification in individual cells.

Whole genome duplication has been observed in many malignancies, including non-small cell lung carcinoma, oesophageal and cervical adenocarcinomas in addition to HGSC, and is frequently, but not exclusively, associated with *TP53* mutations [36]. The timing of WGD during HGSC development remains unclear but our results suggest it is not a late feature. Indeed, the analysis here is consistent with our previous findings in early and late-stage HGSC cohorts [9], and provides evidence that the presence of WGD in STIC may confer a tumour growth advantage and potentially a more aggressive phenotype. Further investigation using *in vivo* models comparing potential proliferative advantages and clonal expansion of p53 mutant clones with and without WGD in the fallopian tube epithelium of mouse models would be informative here.

Although this is one of the few studies to examine the genomics of p53 signatures in detail, our study has several limitations. Firstly, our sample size is small; identifying cases with p53 signatures, STIC lesions and invasive carcinoma in the same specimen requires extensive pathological examination, and the samples require careful and time-intensive laser capture microdissection. Secondly, p53 signature and STIC lesions are microscopic, and can only be identified retrospectively in formalin-fixed, paraffin-embedded samples. The small number of cells in these samples limits the amount of genomic analysis that can be done, especially given the potential for artefacts induced by formalin fixation. Moreover, although we used validated pathological criteria to define lesions [4], demarcations between p53 signatures and STIC lesions are not precise, meaning that samples may be admixed with cells from the immediately adjacent lesion. Third, all our cases had invasive carcinoma meaning that the p53 signatures and STIC lesions had persisted during the development of invasive disease, a process that may take several years [7]. Thus, it is possible that further time-dependent changes had occurred in the pre-invasive disease analysed here. Wang et al very recently described non-random chromosomal alterations in STIC lesions, including those without associated carcinoma, and suggested that there might be two classes of STIC, active and dormant, with greater degrees of aneuploidy in the active group.

They also identified similar copy number gains in matched STIC lesions and invasive carcinoma within individual patients [45]. Nonetheless, our observation that the majority of sequence changes identified in p53 signatures were also present in STIC and other ovarian/metastasis sites supports our evolutionary model and highlights the role of ploidy changes in HGSC evolution. Finally, it is important to note that our analysis primarily focused on shallow whole genome sequencing to obtain ploidy and copy number information, and it was not able to capture allele-specific results for *TP53* and other oncogenes. As stated above, WGS or FISH would allow more detailed analysis of allele-specific changes or make definite statements about WGD, whilst single-cell analyses will allow more comparison of p53 signatures and STIC lesions within the same patient to identify early genomic alterations at much greater resolution.

In summary, our analysis of matched p53 signatures, STIC lesions and invasive carcinoma samples suggests that profound genomic instability is evident very early in the development of high-grade serous carcinoma and that changes suggestive of whole genome duplication are evident in STIC lesions. This suggests that the trajectory of HGSC, and thus patient prognosis, is determined by the point of STIC emergence, highlighting the importance of strategies that will allow earlier detection of HGSC. Moreover, isolated STIC lesions are identified in up to 12% of women undergoing prophylactic salpingectomy or salpingo-oophorectomy [46] and there remains considerable debate as to optimum management as the risk of developing a subsequent invasive high grade serous ovarian or peritoneal carcinoma can be as high as 15% [47]. Our data may help to identify poor prognosis features associated with higher risk of recurrence, and thus guide future clinical management.

## Acknowledgments

This work was funded by an Imperial/China Scholarship Council scholarship to ZC, the NIHR Imperial Biomedical Research Centre (grant number P77646), Ovarian Cancer Action, and Cancer Research UK (grant numbers A15973, A15601, A18072, A17197, A19274 and A19694). IMcN holds an NIHR Senior Investigator Award. OLS is a recipient of grants from La Ligue contre le Cancer, La Fondation Nuovo-Soldati, and Canceropole Lyon Auvergne Rhone-Alpes. Infrastructure support was provided by the Imperial Experimental Cancer Medicine Centre, the Cancer Research UK Convergence Science Centre and the Imperial NIHR Biomedical Research Centre via the Imperial College Healthcare Tissue Bank.

## Authors’ Contributions

Conceptualization: ZC, IAMcN Methodology: ZC, HBM, BL, CB

Investigation: ZC, DPE, BL, HBM, OLS, GR, GG

Pathological examination: CS, BK, NS, JMcD

Clinical samples and data: LAT, JK, IAMcN

Acquisition of funding: IAMcN

Supervision: CB, IAMcN, Writing—original draft: ZC, IAMcN Writing—review & editing: All authors

## Supplemental tables and materials

Supplementary Table 1 Clinical information of STIC patients

Supplementary Table 2 Pathology information of STIC samples

Supplementary Table 3 Purity and ploidy analysis of STIC samples from different pipelines

Supplementary Table 4 Ampliseq result of ovarian and metastasis sites

Supplementary Table 5 Focal copy numbers of COSMIC genes in the node of each patient

Supplementary Figure 1-5 Five representative pathological figures of precursors to high-grade serous carcinoma with staining for p53, Ki67 and H&E. In each patient, the p53 signatures had aberrant p53 expression, normal morphology, but without high proliferative activity (ie low percentage of Ki67 expression). Serous tubal intraepithelial lesions (STILs) are transitional lesions with intermediate morphology findings and proliferative activity between the p53 signature and STIC. Serous tubal intraepithelial carcinoma lesions (STIC) show cytological atypia and loss of polarity on H&E, p53 overexpression due to p53 mutation, and high proliferative activity with strong Ki67 expression.

Supplementary Figure 6 Representative p53, Ki67 and haematoxylin and eosin images from patient STIC_0012. The anatomical sites in the fallopian tube were selected for laser capture microdissection, and this block contained five STIC lesions. In addition, some STIC samples in the adjacent positions were combined to increase the yield to get enough DNA from STIC lesions. P53, Ki67 and H& staining indicated the position of normal fallopian tube epithelia, stroma, p53 signature, STIC, primary tumour1 and 2. Besides, the right ovary was identified primary ovarian tumour. And metastasis carcinoma was found in the omentum and left paracolic peritoneum. The samples in this patient are: 1. STIC1; 2. STIC2+3; 3. STIC4+5; 4. p53 signature; 5. Stroma; 6. Normal fallopian tube epithelium; 7. Primary tumour in right ovary; 8. Metastasis in omentum; 9. Metastasis in left paracolic peritoneum.

Supplementary Figure 7 Representative p53, Ki67 and haematoxylin and eosin images from patient STIC_0014. This block was from the right fallopian tube. Four STIC lesions were combined, and one p53 signature was collected from this block. P53, Ki67 and H&E staining indicated the position of normal fallopian tube epithelium, stroma, p53 signature, STIC (STIC1,2,3,4 together), primary tumour 1 and 2. In addition, the right ovary was identified as the primary ovarian tumour, and metastasis carcinoma was found in the omentum. The samples in this patient are: 1. STIC1+2+3+4; 2. p53 signature; 3. invasive tumour 1; 4. invasive tumour 2; 5. Normal fallopian tube epithelium; 6. Stroma; 7. invasive tumour in right ovary; 8. Metastasis in omentum.

Supplementary Figure 8 Representative p53, Ki67 and haematoxylin and eosin images from patient STIC_0013. This block is from a stage I patient, and there was no metastasis carcinoma in this patient. The anatomical sites in the fallopian tube were selected for laser capture microdissection, and this block contained two p53 signatures and four STIC lesions. Two p53 signatures were combined as a p53 signature sample, and four STIC lesions were combined as two STIC samples based on adjacent locations in the block. P53, Ki67 and H& staining were performed on three slides as an indicator of the position of stroma, p53 signature3+5, STIC1+2, STIC4+6, invasive tumour 7 and 8. The samples in this patient are: 1. p53 signature3+5; 2.STIC2+3; 3.STIC4+6; 4.invasive tumour 7; 5.invasive tumour 8; 6.Stroma.

Supplementary Figure 9 Representative p53, Ki67 and haematoxylin and eosin images from patient STIC_0001. The anatomical sites in the fallopian tube were selected for laser capture microdissection, and this block contained STIL and STIC lesions. P53, Ki67 and H& staining were performed on three slides as an indicator of the position of stroma, STIL, STIC. The samples in this patient are: 1. STIL; 2. STIC; 3. Stroma; 4. primary tumour in left ovary; 5. metastasis in omentum; 6. metastasis tumour in right paracolic gutter; 7. metastasis tumour in aorto-caval lymph node.

Supplementary Figure 10 Comparison of different bin sizes in STIC cohort. (A) number of segments (B) relative errors (C) Purity and (D) Ploidy estimates from ACE package

Supplementary Figure 11 Analysis of copy number alterations phylogeny in STIC cohort. Phylogenetic trees reconstructed by CNETML using relative total copy numbers called from multi-region shallow whole genome sequencing (sWGS) data of STIC cohort. The bootstrap support values are depicted in coloured rectangles, with lighter colours indicating stronger support. The coloured bars at the internal nodes indicate the confidence intervals of node heights. Each internal node in the tree is identified by an ID (left panel), which is also indicated in the heatmap illustrating copy number alterations across the genome in each patient (right panel).

Supplementary Figure 12 The ploidy comparison in missense and non-missense of TP53 mutations in early stage/late stage cohort (A)-(B) (no significance, p>0.05).

## References

1. Piek JM, van Diest PJ, Zweemer RP, et al. Dysplastic changes in prophylactically removed Fallopian tubes of women predisposed to developing ovarian cancer. J Pathol 2001; 195: 451–456.

2. Medeiros F, Muto MG, Lee Y, et al. The tubal fimbria is a preferred site for early adenocarcinoma in women with familial ovarian cancer syndrome. The American journal of surgical pathology 2006; 30: 230–236.

3. Mehra K, Mehrad M, Ning G, et al. STICS, SCOUTs and p53 signatures; a new language for pelvic serous carcinogenesis. Front Biosci (Elite Ed) 2011; 3: 625–634.

4. Vang R, Visvanathan K, Gross A, et al. Validation of an algorithm for the diagnosis of serous tubal intraepithelial carcinoma. International journal of gynecological pathology: official journal of the International Society of Gynecological Pathologists 2012; 31: 243–253.

5. Ducie J, Dao F, Considine M, et al. Molecular analysis of high-grade serous ovarian carcinoma with and without associated serous tubal intra-epithelial carcinoma. Nat Commun 2017; 8: 990.

6. Eckert MA, Pan S, Hernandez KM, et al. Genomics of Ovarian Cancer Progression Reveals Diverse Metastatic Trajectories Including Intraepithelial Metastasis to the Fallopian Tube. Cancer Discov 2016; 6: 1342–1351.

7. Labidi-Galy SI, Papp E, Hallberg D, et al. High grade serous ovarian carcinomas originate in the fallopian tube. Nat Commun 2017; 8: 1093.

8. Smith P, Bradley T, Gavarró LM, et al. The copy number and mutational landscape of recurrent ovarian high-grade serous carcinoma. Nat Commun 2023; 14: 4387.

9. Cheng Z, Mirza H, Ennis DP, et al. The Genomic Landscape of Early-Stage Ovarian High-Grade Serous Carcinoma. Clin Cancer Res 2022; 28: 2911–2922.

10. Li H, Durbin R. Fast and accurate short read alignment with Burrows-Wheeler transform. *Bioinformatics (Oxford*, England) 2009; 25: 1754–1760.

11. Sandmann S, de Graaf AO, Karimi M, et al. Evaluating Variant Calling Tools for Non-Matched Next-Generation Sequencing Data. Scientific reports 2017; 7: 43169.

12. Benjamin D, Sato T, Cibulskis K, et al. Calling Somatic SNVs and Indels with Mutect2. bioRxiv 2019: 861054.

13. Saunders CT, Wong WS, Swamy S, et al. Strelka: accurate somatic small-variant calling from sequenced tumor-normal sample pairs. *Bioinformatics (Oxford*, England) 2012; 28: 1811–1817.

14. Scheinin I, Sie D, Bengtsson H, et al. DNA copy number analysis of fresh and formalin-fixed specimens by shallow whole-genome sequencing with identification and exclusion of problematic regions in the genome assembly. Genome research 2014; 24: 2022–2032.

15. van de Wiel MA, Kim KI, Vosse SJ, et al. CGHcall: calling aberrations for array CGH tumor profiles. *Bioinformatics (Oxford*, England) 2007; 23: 892–894.

16. Aujla S, Aloe C, Vannitamby A, et al. Programmed Death-Ligand 1 Copy Number Loss in NSCLC Associates With Reduced Programmed Death-Ligand 1 Tumor Staining and a Cold Immunophenotype. Journal of thoracic oncology: official publication of the International Association for the Study of Lung Cancer 2022; 17: 675–687.

17. Macintyre G, Goranova T, De Silva D, et al. Copy-number signatures and mutational processes in ovarian carcinoma. Nat Genet 2018; 50: 1262–1270.

18. Steele CD, Abbasi A, Islam SMA, et al. Signatures of copy number alterations in human cancer. Nature 2022.

19. Singh N, Piskorz AM, Bosse T, et al. p53 immunohistochemistry is an accurate surrogate for TP53 mutational analysis in endometrial carcinoma biopsies. J Pathol 2020; 250: 336–345.

20. Poell JB, Mendeville M, Sie D, et al. ACE: absolute copy number estimation from low-coverage whole-genome sequencing data. *Bioinformatics (Oxford*, England) 2019; 35: 2847–2849.

21. Adalsteinsson VA, Ha G, Freeman SS, et al. Scalable whole-exome sequencing of cell-free DNA reveals high concordance with metastatic tumors. Nat Commun 2017; 8: 1324.

22. Sondka Z, Bamford S, Cole CG, et al. The COSMIC Cancer Gene Census: describing genetic dysfunction across all human cancers. Nat Rev Cancer 2018; 18: 696–705.

23. Nagasawa S, Ikeda K, Horie-Inoue K, et al. Systematic Identification of Characteristic Genes of Ovarian Clear Cell Carcinoma Compared with High-Grade Serous Carcinoma Based on RNA-Sequencing. International journal of molecular sciences 2019; 20: 4330.

24. Akahane T, Masuda K, Hirasawa A, et al. TP53 variants in p53 signatures and the clonality of STICs in RRSO samples. Journal of gynecologic oncology 2022; 33: e50.

25. Karst AM, Jones PM, Vena N, et al. Cyclin E1 deregulation occurs early in secretory cell transformation to promote formation of fallopian tube-derived high-grade serous ovarian cancers. Cancer Res 2014; 74: 1141–1152.

26. Bronder D, Tighe A, Wangsa D, et al. TP53 loss initiates chromosomal instability in fallopian tube epithelial cells. Dis Model Mech 2021; 14.

27. Mei J, Tian H, Huang HS, et al. Cellular models of development of ovarian high-grade serous carcinoma: A review of cell of origin and mechanisms of carcinogenesis. Cell Prolif 2021; 54: e13029.

28. Mermel CH, Schumacher SE, Hill B, et al. GISTIC2.0 facilitates sensitive and confident localization of the targets of focal somatic copy-number alteration in human cancers. Genome biology 2011; 12: R41.

29. TCGA. Integrated genomic analyses of ovarian carcinoma. Nature 2011; 474: 609–615.

30. Kim JE, Choi J, Sung CO, et al. High prevalence of TP53 loss and whole-genome doubling in early-onset colorectal cancer. Exp Mol Med 2021; 53: 446–456.

31. Lu B, Curtius K, Graham TA, et al. CNETML: maximum likelihood inference of phylogeny from copy number profiles of multiple samples. Genome biology 2023; 24: 144.

32. Baslan T, Morris JPt, Zhao Z, et al. Ordered and deterministic cancer genome evolution after p53 loss. Nature 2022; 608: 795–802.

33. Ahmed AA, Etemadmoghadam D, Temple J, et al. Driver mutations in TP53 are ubiquitous in high grade serous carcinoma of the ovary. J Pathol 2010; 221: 49–56.

34. Köbel M, Kang EY, Weir A, et al. p53 and ovarian carcinoma survival: an Ovarian Tumor Tissue Analysis consortium study. The journal of pathology Clinical research 2023; 9: 208–222.

35. McGranahan N, Favero F, de Bruin EC, et al. Clonal status of actionable driver events and the timing of mutational processes in cancer evolution. Sci Transl Med 2015; 7: 283ra254.

36. Bielski CM, Zehir A, Penson AV, et al. Genome doubling shapes the evolution and prognosis of advanced cancers. Nat Genet 2018; 50: 1189–1195.

37. Zeng J, Hills SA, Ozono E, et al. Cyclin E-induced replicative stress drives p53-dependent whole-genome duplication. Cell 2023; 186: 528–542.e514.

38. Nichols CA, Gibson WJ, Brown MS, et al. Loss of heterozygosity of essential genes represents a widespread class of potential cancer vulnerabilities. Nat Commun 2020; 11: 2517.

39. Ciriello G, Miller ML, Aksoy BA, et al. Emerging landscape of oncogenic signatures across human cancers. Nat Genet 2013; 45: 1127–1133.

40. Shih IM, Wang Y, Wang TL. The Origin of Ovarian Cancer Species and Precancerous Landscape. Am J Pathol 2021; 191: 26–39.

41. Engqvist H, Parris TZ, Biermann J, et al. Integrative genomics approach identifies molecular features associated with early-stage ovarian carcinoma histotypes. Scientific reports 2020; 10: 7946.

42. Saotome K, Chiyoda T, Aimono E, et al. Clinical implications of next-generation sequencing-based panel tests for malignant ovarian tumors. Cancer medicine 2020; 9: 7407–7417.

43. Murai K, Dentro S, Ong SH, et al. p53 mutation in normal esophagus promotes multiple stages of carcinogenesis but is constrained by clonal competition. Nat Commun 2022; 13: 6206.

44. Knudson AG, Jr. Mutation and cancer: statistical study of retinoblastoma. Proc Natl Acad Sci USA 1971; 68: 820–823.

45. Wang Y, Douville C, Chien Y-W, et al. Aneuploidy Landscape in Precursors of Ovarian Cancer. Clin Cancer Res 2023.

46. Visvanathan K, Shaw P, May BJ, et al. Fallopian Tube Lesions in Women at High Risk for Ovarian Cancer: A Multicenter Study. *Cancer prevention research (Philadelphia*, Pa) 2018; 11: 697–706.

47. Ruel-Laliberté J, Kasasni SM, Oprea D, et al. Outcome and Management of Serous Tubal Intraepithelial Carcinoma Following Opportunistic Salpingectomy: Systematic Review and Meta-Analysis. J Obstet Gynaecol Can 2022; 44: 1174–1180.

